# Data Descriptor: An open-access CT-based 3D anatomical dataset of extant sharks across all major lineages

**DOI:** 10.64898/2026.06.29.734410

**Authors:** Suyuan Yao, Xiaoyue Liu, Yemao Hou, Pengfei Yin, Xiaomei Zhang, Xuejing Cui, Jing Lu

**Affiliations:** Key Laboratory of Vertebrate Evolution and Human Origins, Institute of Vertebrate Paleontology and Paleoanthropology, Chinese Academy of Sciences, Beijing 100044, China; University of Chinese Academy of Sciences, Beijing 100049, China; China University of Geosciences, Wuhan 430074, China

## Abstract

Sharks exhibit extraordinary morphological diversity across a wide range of ecological niches, yet large-scale, high-resolution digital datasets of their internal anatomy remain limited. Here we present an open-access 3D shark anatomical repository derived from published X-ray computed tomography (CT) data, featuring manually segmented and systematically annotated models of the chondrocranium, visceral arches, axial skeleton, musculature, and viscera in standard STL format. The dataset comprises 117 individuals, representing 72 species across 25 families and all nine extant shark orders, with 115 full-body reconstructions and two head-only models. This open-access dataset offers a comprehensive resource for comparative anatomy, biomechanical simulations, evolutionary developmental biology and biomimetics research of extant sharks.

## 1 Background & Summary

Sharks (Selachii) represent one of the most globally distributed and morphologically diverse radiations of jawed vertebrates^1–3^. Their skeletal and soft-tissue systems display extensive structural diversity associated with divergent feeding strategies^4–6^, locomotion^7,8^, and ecological adaptation^9,10^. Detailed, high-quality anatomical data are therefore crucial for investigating functional morphology, evolutionary developmental biology, and macroevolutionary patterns in sharks^9,11,12^.

The advent of X-ray computed tomography (CT) has transformed vertebrate morphological research by enabling the non-destructive, high-resolution visualization of complicated skeletal and soft-tissue anatomy^13,14^. CT-based three-dimensional (3D) reconstruction has become an essential tool in comparative anatomy, geometric morphometrics, and biomechanical modelling in chondrichthyan fishes^15–18^. However, previous research has mainly focused on species-specific^16,19^ and isolated anatomical structures, such as cranial morphology^17^, jaw mechanics^5,20^ or oral denticles^21–23^, leaving comprehensive, whole-body 3D morphological datasets of sharks remarkably scarce.

To address this gap, we present an open-access, fully segmented 3D anatomical dataset dedicated to extant sharks, based on validated and previously published CT data^24^. Comprising 117 individuals of 72 species, our dataset encompasses 27 families (covering ∼79% of the 34 recognized shark families) and 44 genera (∼42% of 105 genera) across all nine extant orders (Fig.1). Compared with existing chondrichthyan digital resources, such as the Chondrichthyan Tree of Life (CTOL) project, our dataset offers significantly deeper taxonomic resolution within Selachii, capturing 10 additional families, 14 additional genera, and 58 previously unreconstructed species. This includes complete anatomical reconstructions for deep-water and historically underrepresented lineages, such as Hexanchiformes, Pristiophoriformes, and Squatiniformes.

**Figure 1.**
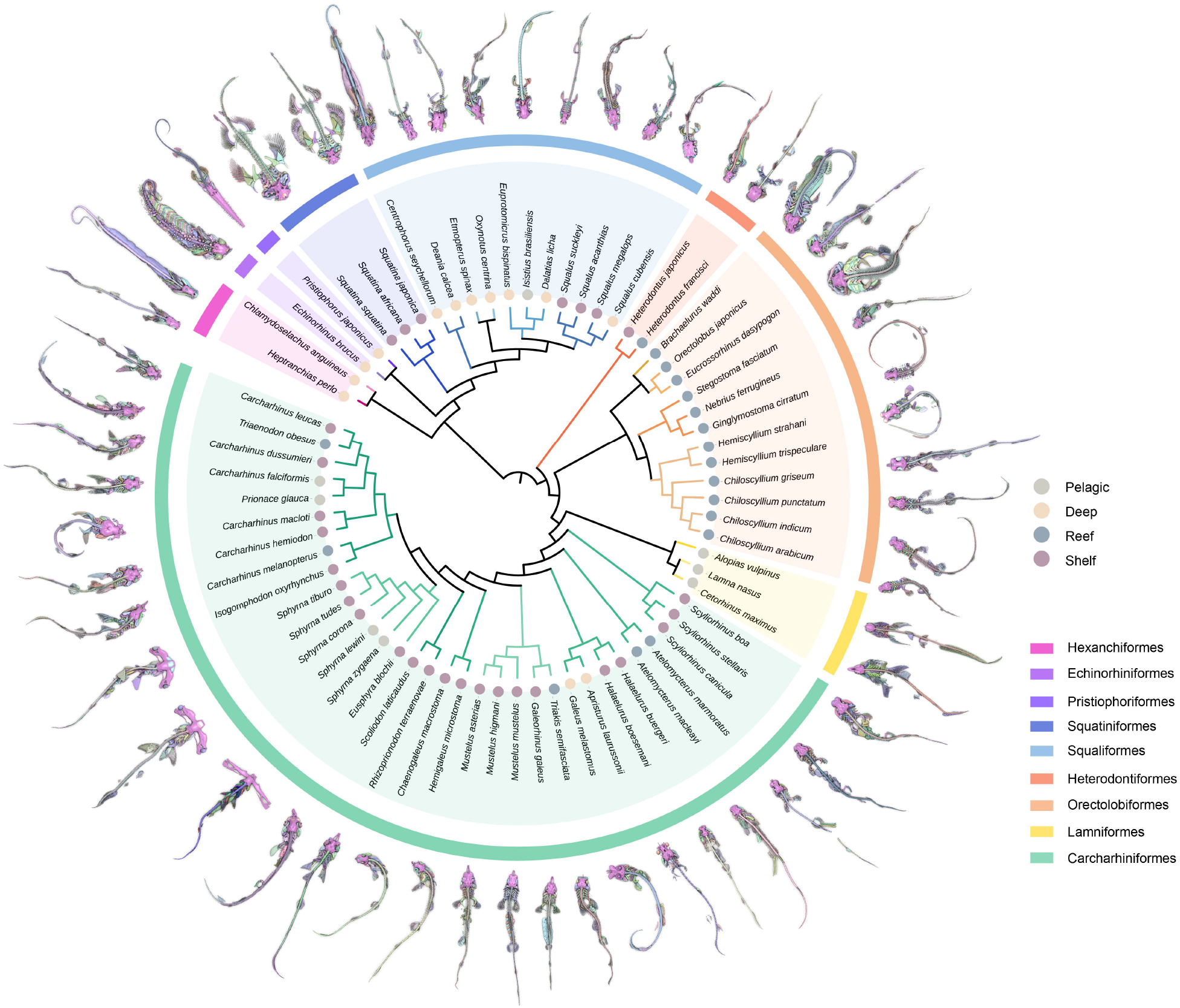
Phylogenetic framework illustrating the taxonomic coverage of the 3D anatomical shark dataset. Pruned circular phylogeny derived from the VertLife Project shark tree^3^ (10cal.tree; http://vertlife.org/sharktree/), showing the taxonomic coverage across nine extant orders (coloured ring legend). Each terminal branch is annotated with a fully segmented 3D skeletal model of the representative species, alongside habitat classifications (Pelagic, Deep, Reef, Shelf).

Unlike traditional datasets that focus on isolated skeletal elements^25^, our models integrate multiple anatomical modules, including the chondrocranium, visceral arches, axial and appendicular skeleton, musculature, and visceral organs. Supported by uniform segmentation protocols and standard metadata across all specimens, these models are optimized for quantitative analytical workflows, including geometric morphometrics (GMM), principal component analyses (PCA), and computational biomechanical simulations. By expanding taxonomic coverage within Selachii and providing detailed three-dimensional reconstructions, this dataset complements existing chondrichthyan resources and facilitates high-resolution investigations of shark functional morphology and evolutionary research.

## 2 Methods

### 2.1 Specimens and CT data acquisition

The 3D reconstructions in this study are based on an open-access CT repository of 122 individuals representing 73 extant shark species, originally established by Kamminga et al. (2017) and hosted on Figshare (Data Citation 1). The source material consists of specimens primarily housed in the spirit collections of the Naturalis Biodiversity Center (NBC) and the British Museum of Natural History (BMNH). Scanning was conducted using optimized protocols on two medical platforms: NBC material was acquired at the Leiden University Medical Center, the Netherlands (LUMC) via a Toshiba Aquilion 64 (100 kV, 150 mAs, 0.5 mm slice thickness), while BMNH material was processed at the Royal Brompton and Harefield NHS Trust (RBH) using a Siemens Somatom Sensation 64 (100 kV, 210 mAs, 1.0 mm slice thickness)^24^.

To ensure high anatomical accuracy and consistency of the segmented models, each CT image stack was evaluated to determine its suitability for 3D reconstruction. From the initial collection of 122 specimens, a total of 117 individuals (representing 72 species) met the predefined quality criteria and were retained for subsequent 3D reconstruction and segmentation. The final dataset comprises 115 full-body and two head-only models, collectively providing a phylogenetically representative digital library of extant shark anatomy.

### 2.2 Digital segmentation and 3D reconstruction

Raw CT scan data were imported into Mimics, v 19.0 (Materialise Software) for digital segmentation and three-dimensional reconstruction (Fig.2). Given the limited contrast of medical CT in resolving cartilaginous and soft tissues, a predominantly manual segmentation workflow was employed. Initial segmentation was performed using thresholding based on grey values to isolate high-density skeletal structure.

**Figure 2.**
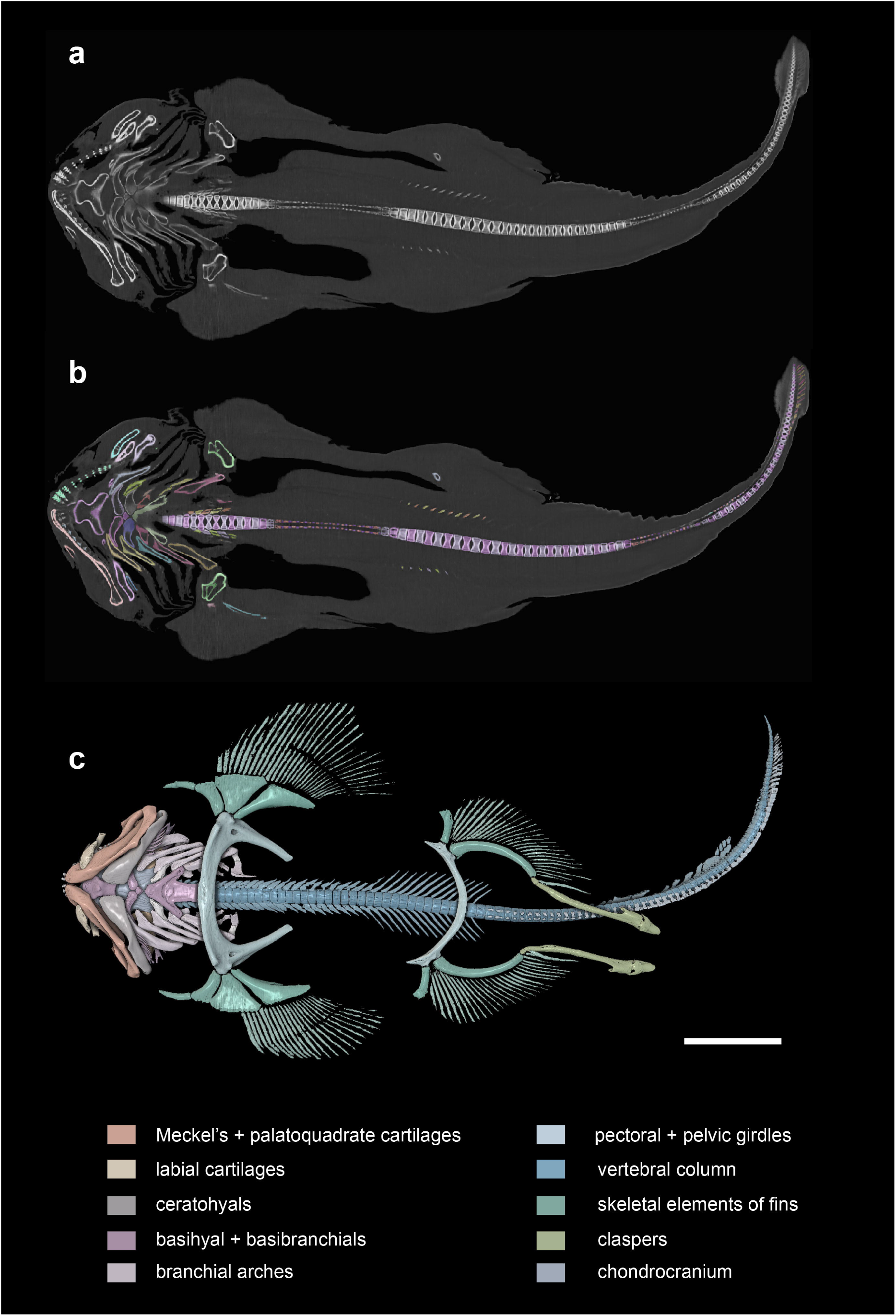
Workflow for three-dimensional (3D) anatomical segmentation and volume rendering of a male *Squatina squatina* (RMNH.PISC.24047). **(a)** Representative two-dimensional tomographic slice from the raw CT dataset visualized in Mimics, showing the initial contrast and skeletal resolution. **(b)** Manual segmentation and refinement of discrete anatomical masks within the same tomographic plane, where distinct colours represent individual segmented components. **(c)** Integrated 3D anatomical model rendered in Vayu, displaying the spatial relationship among major skeletal elements. The colour-coded legend identifies anatomical modules, including the chondrocranium, visceral arches, and appendicular skeleton, etc. Scale bar = 10 cm.

Semi-automatic tools, including region-growing algorithms, were subsequently applied to expand selected regions. These preliminary results were further refined on a slice-by-slice basis using manual editing (e.g., lasso and brush tools) to ensure anatomical accuracy. Each specimen was segmented into multiple anatomical components to generate a comprehensive digital anatomical reconstruction, including chondrocranium, visceral arches, axial and appendicular skeleton, etc.

### 2.3 Anatomical annotation and labeling

Following the segmentation and reconstruction, each segmented structure was exported as an individual three-dimensional surface mesh in STL format, enabling subsequent anatomical annotation and visualization. The level of anatomical segmentation varied among species according to morphological complexity, with the number of segmented elements ranging from 22 to 224 per specimen (mean = 78 elements per specimen), as summarized in the metadata (see Supplementary Table 1). These elements can be grouped into five major functional categories as follows:

#### Craniofacial Skeleton

Chondrocranium, rostral cartilage, labial cartilages. Sensory structures, including the eyeballs and inner ear, were also segmented and reconstructed separately.

#### Visceral Arches

The mandibular arch (Meckel’s cartilages, palatoquadrate cartilages and teeth), the hyoid arch (hyomandibulae, ceratohyals, and basihyal) and the complete branchial series (pharyngobranchials, epibranchials, ceratobranchials, and associated gill rays) were constructed (Fig.3).

**Figure 3.**
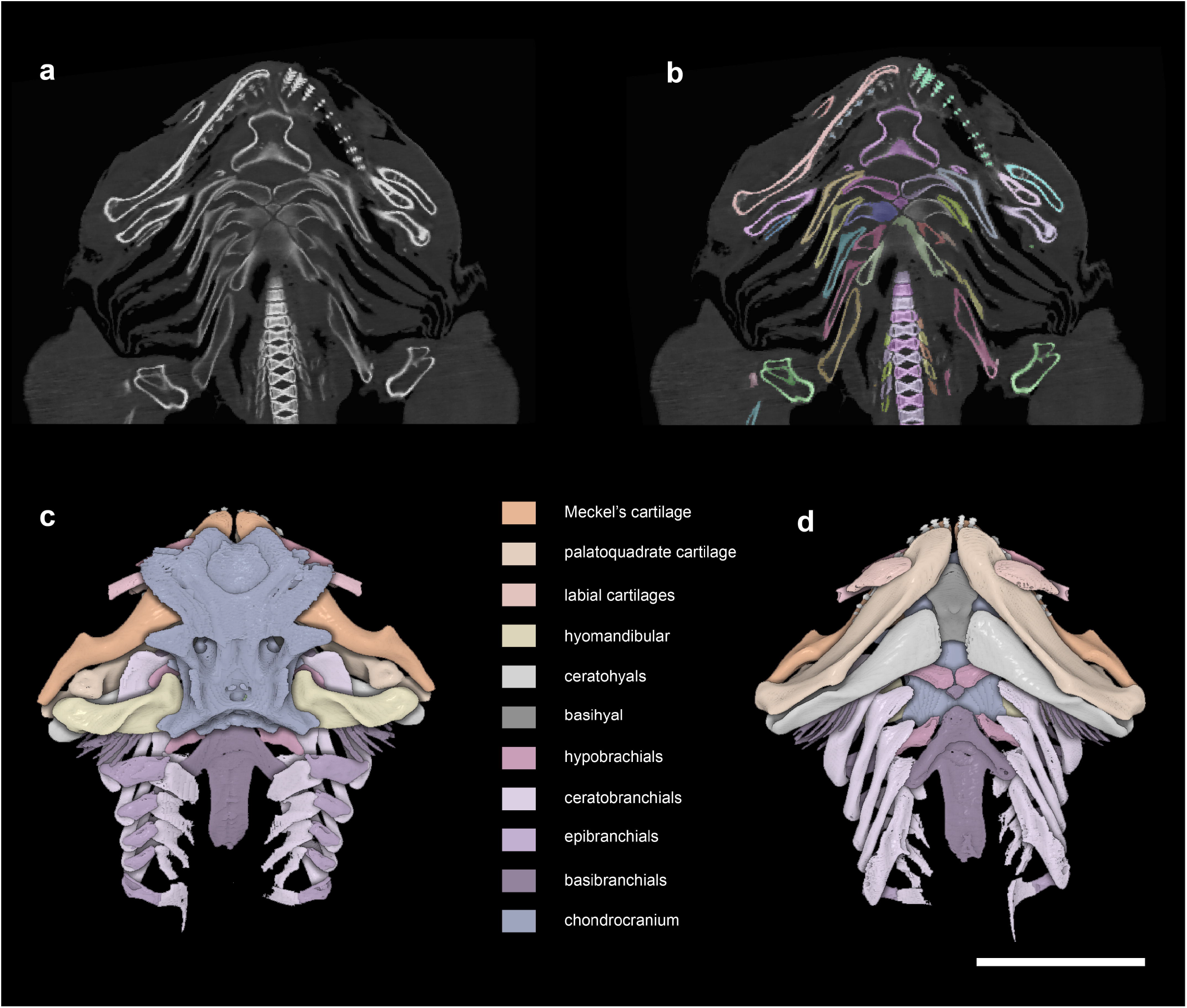
High-resolution 3D reconstruction and anatomical segmentation of the cranial region of a male *Squatina squatina* (RMNH.PISC.24047). **(a)** Representative two-dimensional tomographic slice from the raw CT dataset visualized in Mimics, showing the initial contrast and skeletal resolution. **(b)** Manual segmentation and refinement of discrete anatomical masks within the same tomographic plane, where distinct colours represent individual segmented components. **(c, d)** Three-dimensional reconstructions of the head in dorsal (c) and ventral (d) views, highlighting the chondrocranium and the splanchnocranium, including the mandibular, hyoid, and branchial arches. The colour-coded legend identifies the specific modules for comparative anatomy analyses. Scale bar = 10 cm.

#### Axial and Appendicular Skeleton

The vertebral column (including neural and hemal arches), pectoral and pelvic girdles (with scapular processes), and the full suite of fin skeletal elements (propterygium, mesopterygium, metapterygium, radial pterygiophores, and ceratotrichia).

#### Musculature and Integument

Systematic segmentation of the major cranial and trunk musculature, together with the external dermal envelope (skin). Notably, in a male *Echinorhinus brucus* (BMNH 1900.11.6.7), nearly a hundred individual muscles were reconstructed, providing a detailed reference for future anatomical and functional analyses.

#### Viscera and reproductive structures

Where tissue preservation permitted, major digestive and metabolic organs (including stomach, liver and valvular intestine) were reconstructed. Reproductive structures such as the testis, ovary and embryos were also included where clearly identifiable in the original tomographic scans.

## 3 Data Records

The annotated 3D reconstruction dataset for this study is publicly accessible on the Archives of Digital Morph (ADMorph) database (Data Citation 2), which offers an interactive platform for visualizing and directly downloading individual anatomical components. Each specimen is assigned a unique identifier within the database, while corresponding DOIs and direct download links are provided in the comprehensive metadata (see Supplementary Table 1).

In total, the dataset comprises high-resolution 3D anatomical models for 117 shark specimens, representing 72 extant species from 27 families and covering all nine taxonomic orders (Fig.1). The collection includes 115 complete full-body reconstructions and two head-only models, all provided as standard surface meshes in STL format. For each specimen, discrete anatomical structures (e.g., chondrocranium, branchial arches, pectoral girdles, ceratotrichia) are stored as individual files, allowing users to selectively download targeted components for use in comparative anatomical research (Fig.4).

**Figure 4.**
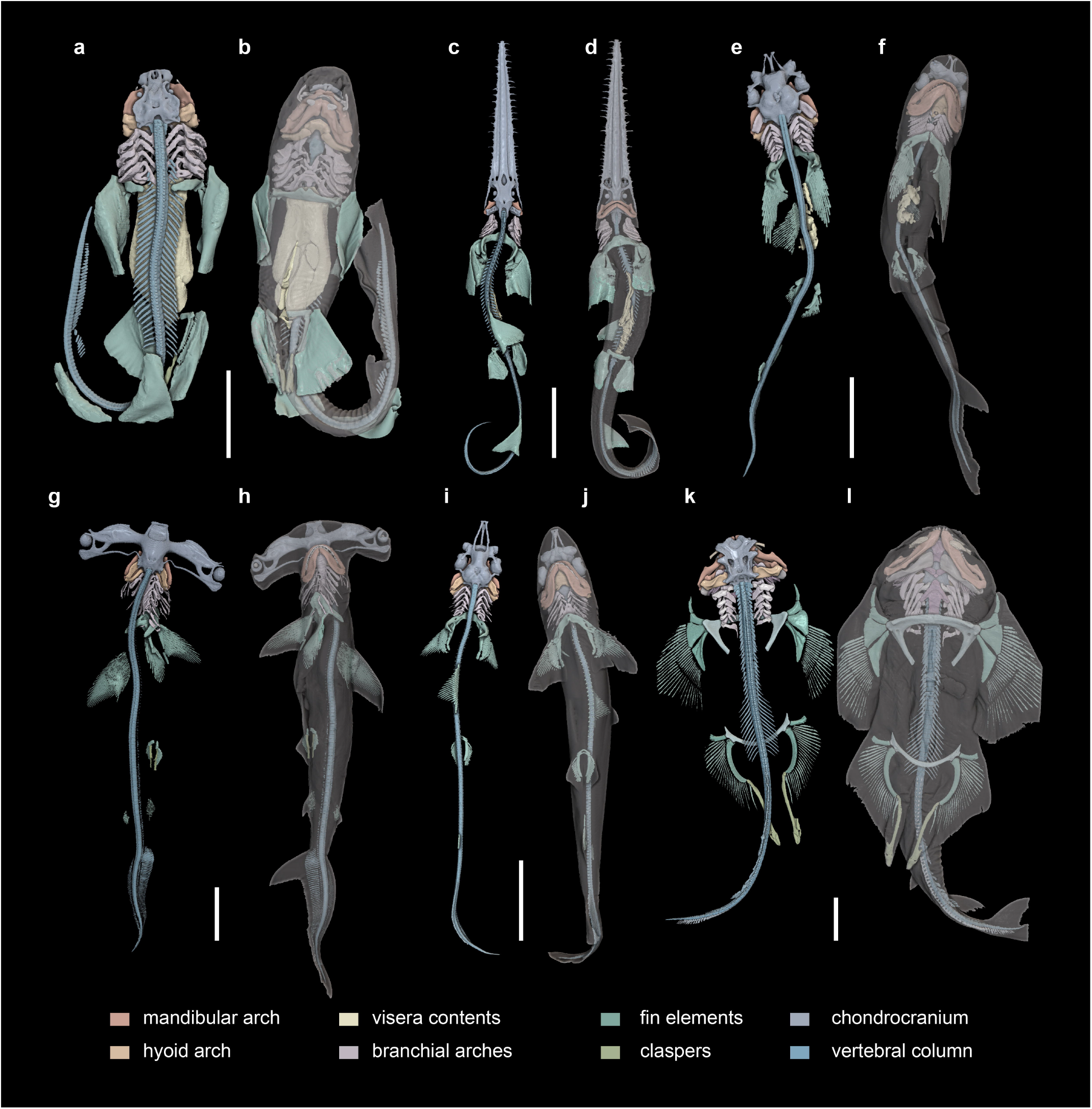
Three-dimensional volume-rendered reconstructions of selected shark specimens, illustrating morphological diversity across taxa. Specimens are shown in dorsal (left) and ventral (right) views with semi-transparent external tissues to reveal internal anatomy. Scale bar = 10 cm. **(a**,**b)** a male *Brachaelurus waddi* (BMNH 1890.9.23.233) **(c**,**d)** a female *Pristiophorus japonicus* (BMNH 1936.7.29.12) **(e**,**f)** a male *Carcharhinus melanopterus* (ZMA.PISC.108682) **(g**,**h)** a female *Sphyrna zygaena* (ZMA.PISC.108725) **(i, j)** a female *Rhizoprionodon terraenovae* (ZMA.PISC.111362) **(k**,**l)** a male *Squatina squatina* (RMNH.PISC.24047)

## 4 Technical Validation

Digital segmentation and three-dimensional (3D) reconstruction were performed using Materialise Mimics (version 19.0) on a set of high-performance workstations (Dell T7920 and Dell Precision 3680) equipped with Intel Xeon and Intel Core i7 processors, 64–192 GB of RAM, and NVIDIA GPUs. The anatomical accuracy of the reconstructed models was primarily determined by the resolution and contrast quality of the source CT data. Segmentation parameters were adjusted and optimized throughout each image stack to accommodate subtle variations in tissue density. All segmented structures were visually inspected and manually refined in orthogonal planes (axial, sagittal, and coronal) to ensure morphological completeness and accuracy.

The segmented models were further examined in Vayu^26^ for visualization and annotation, where distinct colour coding was applied to individual skeletal and tissue structures to improve anatomical clarity. Anatomical annotation and labelling were carried out by a trained research team with expertise in shark morphology, following established anatomical references^27^. This standard technical workflow ensured the production of high-accuracy 3D models suitable for comparative anatomy and educational purposes.

## 5 Usage Notes

The 3D models in this dataset are provided in standard STL format and are freely available for download via the ADMorph database (Data Citation 2) for academic research use. The models are compatible with common software suites (e.g., MeshLab, 3D Viewer, Blender and Vayu) for general visualization, mesh inspection, and high-quality rendering. Individual anatomical structures can be imported independently or simultaneously to reconstruct complete specimen morphology. The ADMorph database also offers a web-based interface for interactive real⍰time anatomical exploration, with no local software installation required.

This dataset enables high-precision quantitative investigations in comparative anatomy, functional morphology and biomechanics. For instance, the high-resolution reconstructions of the Meckel’s and palatoquadrate cartilages support geometric morphometrics and finite element analysis (FEA) of bite force and feeding mechanics^5^. While the intact body surface mesh facilitates computational fluid dynamics (CFD) assessments of hydrodynamic performance and swimming efficiency across diverse shark morphotypes. Note that researchers intending to perform volumetric meshing for FEA or CFD may employ targeted remeshing in software such as ZBrush, Rhinoceros, Geomagic or Autodesk Maya to ensure specific numerical stability requirements and topological integrity.

High-quality renderings of these shark digital models also support science education and public outreach, particularly for museum displays and interactive exhibitions (see Fig.5 and Supplementary video), including immersive virtual reality (VR) experiences and anatomically accurate 3D-printed replicas for teaching.

**Figure 5.**
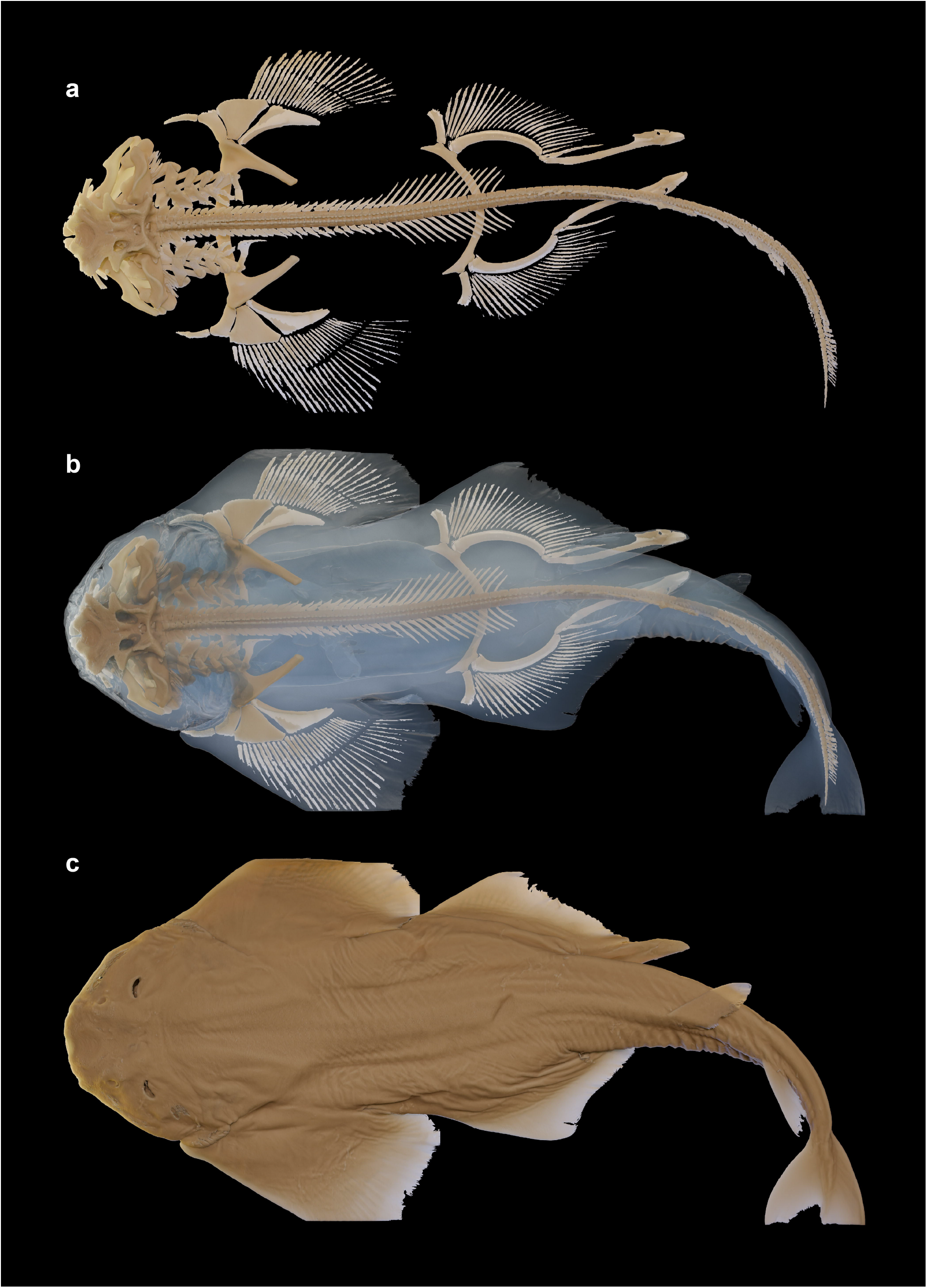
Three-dimensional anatomical visualization of a male *Squatina squatina* (RMNH.PISC.24047), generated using Blender. **(a)** Textured surface rendering of the whole specimen in dorsal view. **(b)** Semi-transparent surface rendering revealing underlying skeletal anatomy. **(c)** Isolated skeletal reconstruction showing the complete axial and appendicular skeleton.

## 6 Data availability

The original CT scan data are deposited on Figshare (Data Citation 1). All 3D models generated in this study are available at the ADMorph database (Data Citation 2). The digital data in standard STL format are publicly accessible via the direct link: http://admorph.ivpp.ac.cn/dataset/specialTopic/detail/insider/22e303fff3bfdf1d9641a72eb2278935 (Note that this link is temporary, as the ADMorph database is currently being optimized for improved stability and long-term accessibility). The data are also permanently deposited on Figshare: https://doi.org/10.6084/m9.figshare.31989363. Complete metadata supporting this study are provided in Supplementary Table 1.

## 7 Code availability

No custom code was used in the preparation of this dataset.

## 9 Acknowledgements

This work was supported by The National Key Research and Development Program of China, Grant No: 2025YFF0811702. We are grateful to X.F. Zhang, C.X. Wang, N. Zhang, and R. Tang for segmenting, three-dimensional rendering and animation production. We acknowledge NICE PaleoVislab, IVPP for biological reconstruction, realistic rendering, and visualization.

## 10. Author information

### Authors contributions

J.L. conceived and supervised the project. J.L. S.Y., and X.L. analyzed, segmented, annotated and revisualized the specimens. J.L. Y.H., P.Y., X.Z. developed ADMorph database. S.Y., X.L. and J.L. discussed and drafted the manuscript. S.Y., J.L. and X.L. generated all figures. S.Y., X.L., Y.H., P.Y., and X.Z. prepared the metadata. All authors discussed and wrote the manuscript.

## Supplementary information

**Supplementary Table 1:** Metadata of shark specimens included in the 3D anatomical dataset.

**Supplementary Video:** Animated 3D visualization of shark specimens displaying the internal anatomical structures through volume-rendered reconstructions generated in Vayu^26^.

